# *in vitro* analysis of a competitive inhibition model for T7 RNA polymerase biosensors

**DOI:** 10.1101/2024.09.23.614532

**Authors:** Ryan Delaney, Katherine A. Lamb, Olivia M. Irvin, Zachary T. Baumer, Timothy A. Whitehead

## Abstract

T7 RNA polymerase (T7 RNAP) biosensors, in which T7 RNAP transcribes some reporter gene or signal in response to external stimuli, have wide applications in synthetic biology and metabolic engineering. We adapted a biochemical reaction network model and used an *in vitro* transcription assay to determine network parameters for different T7 RNAP constructs. Under conditions where template DNA is limiting, the EC_50_ values of native and engineered T7 RNAPs ranged from 33 nM (29 -37 95% c.i.) to 570 nM (258 -714 95% c.i.) (wild-type T7 RNAP). The measured EC_50_ values were largely insensitive to free magnesium, pH, or other buffer conditions. Many biosensor configurations use a split RNAP construct, where the C-terminal (CT7) and N-terminal T7 (NT7) are fused to proximity induced dimerization modules. We used proteolysis and ion exchange chromatography to prepare a CT7 (80 kDa) product. The impact of free CT7 on T7 RNAP transcriptional activity was well described by a competitive inhibition model, with an inhibitory constant K_I_ = 24 nM (22-26 95% c.i.) of the sensor. These model parameters will be useful for forward modeling and design of T7 RNAP-based genetic circuits.

**Highlights:**

- EC_50_ values for natural and engineered T7 RNAPs measured in vitro
- C-terminal T7 RNAP is a competitive inhibitor of full-length T7 RNAP
- The inhibition constant of C-terminal T7 RNAP is lower than the EC_50_ of T7 RNAP

## Introduction

Single subunit RNA polymerases like T7 (T7 RNAP) have become enabling tools in synthetic biology and metabolic engineering due to their relatively simple mechanism of transcription. Natural and engineered versions of T7 RNAP have applications ranging from *in vitro* transcription of mRNA vaccines_**1**_, recombinant protein production^2^, and instantiation of novel genetic circuits ^3–8^. One major use case for T7 RNAP are as biosensor transducers using either cell-free systems or living cells ^9–16^. Many such genetically encoded biosensors use a configuration where T7 RNAP is split into two or more individual genes. These split proteins are genetically encoded as fusions to proteins which come into proximity upon binding a chemical or biomolecule ^17^. Proximity of the split T7 RNAP pieces reconstitute functional polymerase. In this way, chemical inputs can be transduced by T7 RNAP into a transcriptional output.

While these split T7 RNAP sensors can be engineered for high transcriptional fold changes in the presence of ligand, the sensitivities and EC_50_ values are rarely reported. When reported ^9,15^, the EC_50_ values are in the μM range even though the chemically induced proximity partners sense in the high pM-low nM range ^18–20^. T7 RNAP constructs are often split as an N-terminal (NT7) and a ∼80 kDa C-terminal (CT7) piece near amino acid position 179/180 which maintains the promoter specificity loop responsible for binding promoter DNA. CT7 prepared by proteolytic cleavage of full-length T7 RNAP maintains the ability to bind promoter DNA^21,22^. Based on this biochemistry, we hypothesize that free CT7 may competitively inhibit transcription by reconstituted T7 RNAP, diminishing the sensitivity of the biosensor.

In this work, we adapted a biochemical network of T7 RNAP activity that includes the impact of free CT7 on transcription. We then investigated this model, under conditions of excess polymerase, using an *in vitro* transcriptional aptamer assay. We found that both the EC_50_ and activity differ between natural and engineered T7 RNAPs by up to an order of magnitude. We prepared CT7 and showed that it acts as a competitive inhibitor of T7 RNAP transcription, with a measured inhibitory constant in the low to mid nM. These results offer insight into how more sensitive T7 RNAP biosensors can be designed, and will help predictive modeling of T7 RNAP genetic circuits in cell-free and living cell modalities.

## Material and Methods

### Protein expression and purification

Plasmids pET-Express-T7-His6 (WT; pZB017) and pET-Express-T7-His6-TS (TS; pZB030) were from Baumer et al^7^. Both encode a T7 RNAP variant with an N-terminal His_6_ tag. T7 RNAP^TS^ has the following mutations: S430P, N433T, S633P, F849I, F880Y^23^. We also cloned a thermally stabilized N-29-1^TS^ T7 RNAP (L32S, E35G, K98R, Q107K, T122S, A144T, S430P, N433T, S633P, F849I, F880Y)^9^ into the same expression plasmid, forming pZB666. This plasmid was sequence confirmed by Oxford nanopore (Plasmidsaurus).

All full-length RNAPs were expressed and purified according to Baumer et al^7^. CT7 preparation was performed by the addition of trypsin to 3 mg/mL N-29-1^TS^ with a mass ratio of 750:1 (N-29-1^TS^:trypsin) for 2 hours at room temperature in 50mM Tris, 50mM NaCl, 1mM EDTA, 5mM 2-Mercaptoethanol, 7.5 (w/v)% glycerol pH 7.9. The digestion was quenched with a 10-fold molar excess soybean trypsin inhibitor (Sigma Aldrich Cat #10109886001) for 10 minutes. CT7 was separated from the reaction mixture using P11 phosphocellulose (Whatman) resin equilibrated with 50mM sodium phosphate 20mM NaCl pH 7.5 and eluted at 500mM NaCl, 50mM sodium phosphate pH 7.5. CT7 eluate was buffer exchanged using a 30kDa Amicon filter into 50mM tris, 50mM NaCl,, 1mM EDTA, 5mM 2-Mercaptoethanol, 7.5 (w/v)% glycerol pH 7.9 for storage at 4^°^C. Protein concentrations were estimated using predicted A_280_ extinction coefficients (WT: 138830 M^-1^cm^-1^, TS: 140110 M^-1^cm^-1^, N-29-1^TS^: 140110 M^-1^cm^-1^, CT7: 129300 M^-1^cm^-1^), normalized by purity. Purity was assessed by gel densitometry using a denaturing SDS-PAGE Gel with 5 μg of total loaded protein.

### Biochemical assays

Except where noted, fluorescent aptamer assays were performed as described^7^ using the following assay conditions with varying concentration of T7 RNAP variant and CT7, 3nM template DNA containing pT7 promoter (−17:-1 in bold **TAATACGACTCACTATA**GGGAG) and DNA encoding the RNA Peppers aptamer, 16mM NTPs (Thermo cat #R0481), and 1 μM4-Cyano-α-[[4-[(2-hydroxyethyl)methylamino]phenyl]methylene]benzeneacetonitrile (HBC530)^24^. The final reaction buffer contained 128 mM HEPES, 22.5 mM MgCl_2_, 40mM DTT, 1.28mM spermidine, 50mM KCl, 0.1 mg/ml BSA, and 0.001 U/ml Inorganic Pyrophosphatase at pH 7.5.

## Theory and Calculations

We formulated a biochemical reaction network model for T7 RNAP, and T7 RNAP inhibition by CT7, according to Chamberlin and Ring^25^ with modifications noted. T7 RNAP (R) binds its promoter (P) reversibly as an initiation complex (RP_i_). This initiation complex can reversibly form the active elongation complex (RPe), which then leaves the promoter region at a characteristic rate.

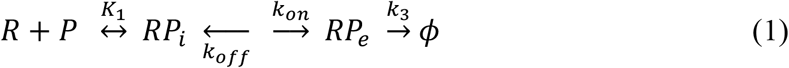

In our model, CT7 competes with the promoter to form an inactive complex I:

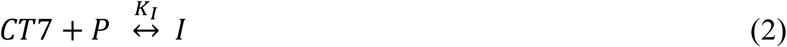

Under steady state conditions, the concentration of the active elongation complex is proportional to activity (amount of transcript produced per unit time), holding NTP concentrations and other reaction conditions constant. k is the proportionality constant in this model:

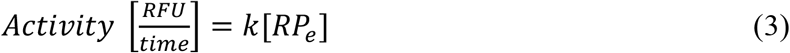

At steady state, we can define K_2_ as the ratio of RP_e_ to RPi:

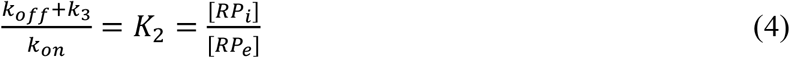

Under conditions where the amount of T7 RNAP is in great excess to promoter DNA, we can solve the mass balance on promoter, where total promoter concentration is P_T_ and total RNA Polymerase concentration is R_T_.

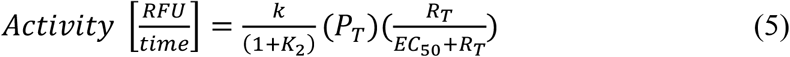

Where EC_50_ is defined as:

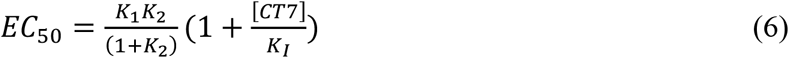

## Results

To evaluate the ability of the model to inform T7 RNAP activity, we used a previously described *in vitro* transcriptional assay. In this assay, template dsDNA containing a T7 promoter and DNA encoding a peppers RNA aptamer is transcribed by T7 RNAP. The peppers aptamer complexes with the chromophore HBC530, allowing for a linear relationship between fluorescence rate (RFU/time) and transcriptional activity (**Fig 1a**). We characterized assay conditions and confirmed that excess NTPs and >5 mM free Mg^2+^ allow for maximal transcriptional activity^25^. Under conditions where the thermally stabilized T7 RNAP^TS^ is in large excess compared with the DNA template, activity varied linearly between 0.2-20 nM dsDNA (**Fig 1b**), as expected from the biochemical model (eqn. 5). The activity was nearly identical with 1 μM and 20 μM HBC530. Therefore, we used 1 μM HBC530 and 3 nM dsDNA template for all further assays.

**Figure 1.**
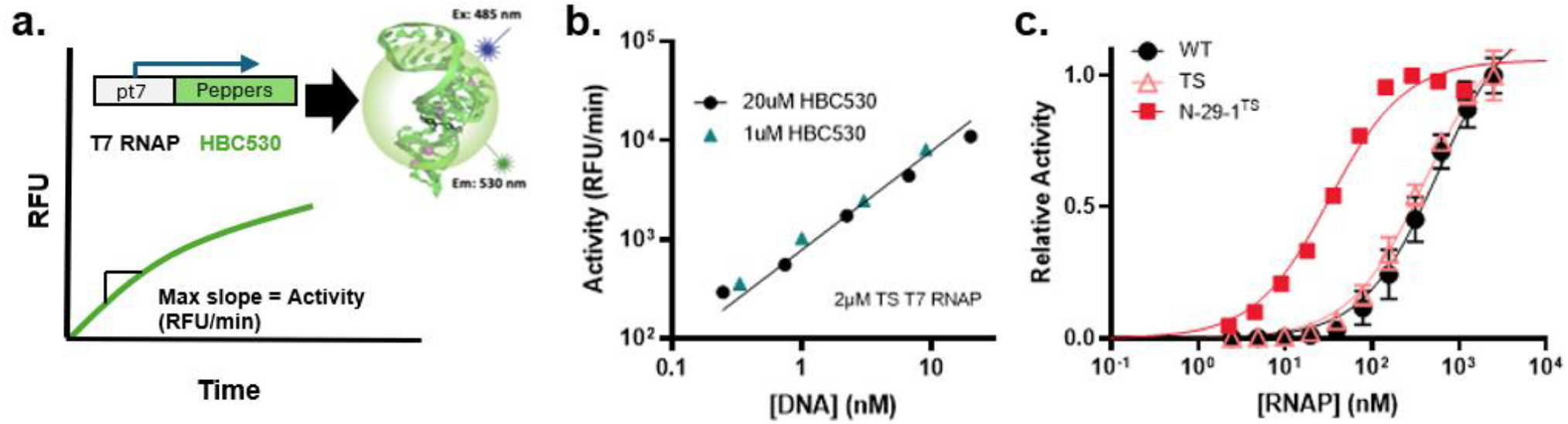
in vitro transcriptional assay for assessing T7 RNAP biochemical parameters. a. Assay design. T7 RNAP is incubated at 37oC with 16mM rNTPs, HBC530, and a dsDNA template containing a canonical T7 promoter sequence and the Peppers gene. Transcription drives formation of the Peppers aptamer, with activity defined as the maximum relative fluorescence units (RFU) over time for the fluorescent Peppers-HBC530 complex. b. Activity (RFU/min) vs. template DNA concentration [DNA] in the presence of 2 μM T7 RNAP and at indicated concentrations of HBC530. In this regime, the transcriptional activity is linear with respect to DNA concentration. The best-fit linear regression line for the 20 μM HBC530 data is shown as a black line (R2=0.98). c. Relative activity vs. indicated T7 RNAP variant [RNAP] using 3 nM template DNA and 1 μM HBC530. For panels b. and c., the error bars represent 1 s.d. of at least three technical replicates. In some cases, the error bars are smaller than the symbol.

Titrations of T7 RNAP (‘WT’) and a thermally stabilized T7 RNAP^TS^ (‘TS’) resulted in saturable relative activity isotherms well fit by an EC_50_ of 570 nM (258 -714 95% c.i.) and 405 (346 -477 95% c.i.), respectively (**Figure 1c, Table 1**). We noticed a small amount (<2-fold) of day-to-day variability in the EC_50_ values, depending on the protein prep (**Table 1**). We also produced and purified a thermally stabilized variant of the N-29-1 T7 RNAP (N-29-1^TS^) originally described by Dickinson et al^9^. The EC_50_ of this engineered polymerase was 33 nM (29 -37 95% c.i.), over an order of magnitude lower than that of WT. Notably, N-29-1^TS^ also had a more than 5-fold reduction in maximum activity than WT (**Figure 1c, Table 1**). While the EC_50_ values we report are broadly consistent with *in vitro* results obtained in the regime of limiting DNA^26^, they are much larger than values reported in assays where T7 RNAP is limiting and template DNA is titrated^27^. To test whether our EC_50_ values reported depend on specific assay conditions, we tested the effect of free magnesium and buffer on the responsiveness of TS. There were small changes in EC_50_ values between conditions, with the lowest magnesium concentration of 13.5 mM yielding a 120 nM EC_50_ for the TS construct (**Table 1**).

**Table 1.** EC_50_, inhibitory constants, and relative activity of natural and engineered T7 RNAPs.

| T7 RNAP | Buffer Conditions | EC<sub>50</sub> [nM] | Relative Activity<sup>‡</sup> |
| --- | --- | --- | --- |
| WT | 128 mM HEPES, 22.5 mM MgCl<sub>2</sub> pH 7.5 | 570<br>(95% c.i.: 258 - 714) | 1.85 (1.7 - 2.0) |
| TS | 128 mM HEPES, 22.5 mM MgCl<sub>2</sub> pH 7.5 | 405<br>(95% c.i.: 346 - 477) (rep 1)<br>236<br>(95% c.i.: 183 - 304) (rep 2) | 1 |
|  | 102 mM HEPES, 18 mM MgCl<sub>2</sub> pH 7.5 | 197<br>(95% c.i.: 136 - 285) | 1.1 |
|  | 76.8 mM HEPES, 13.5 mM MgCl<sub>2</sub> pH 7.5 | 120<br>(95% c.i.: 100-140) | 0.5 |
|  | 30mM Tris, 20mM MgCl<sub>2</sub> pH 7.9 | 219<br>(195 - 245 95% c.i.) | 1.4 |
| N-29-1<sup>TS</sup> | 128 mM HEPES, 22.5 mM MgCl<sub>2</sub> pH 7.5 | 33<br>(95% c.i.: 29 - 37) (rep 1) | 0.4<br>(0.35 - 0.45) (rep 1) |
|  |  | 28<br>(95% c.i.: 24 - 32) (rep 2) | 0.4<br>(0.39 - 0.41) (rep 2) |

<sup>‡</sup> Raw activities (RFU/min) are normalized to 1.0 for TS T7 RNAP under 28 mM HEPES, 22.5 mM MgCl<sub>2</sub> pH 7.5.

To determine whether free CT7 inhibits transcription, we tested two expression constructs of CT7 fused to PYR1 biosensors^19^ with and without a maltose binding protein solubilization tag. Neither construct resulted in soluble expression of pure CT7 (*data not shown*). As an alternative, we prepared CT7 using a trypsin digestion of full-length T7 RNAP (**Fig 2a**), followed by selective adsorption of the DNA binding portion of the C terminal RNAP onto a cation exchange resin. We screened several different T7 RNAP constructs and found that, over the span of two hours, N-29-1^TS^ was fully degraded into a primarily 80 kDa CT7 piece (**Fig 2a**). Selective adsorption and elution from the cation exchange resin resulted in a reasonably pure CT7 (**Fig 2b**) with no transcriptional activity in the aptamer assay (**Fig 2c**).

**Figure 2.**
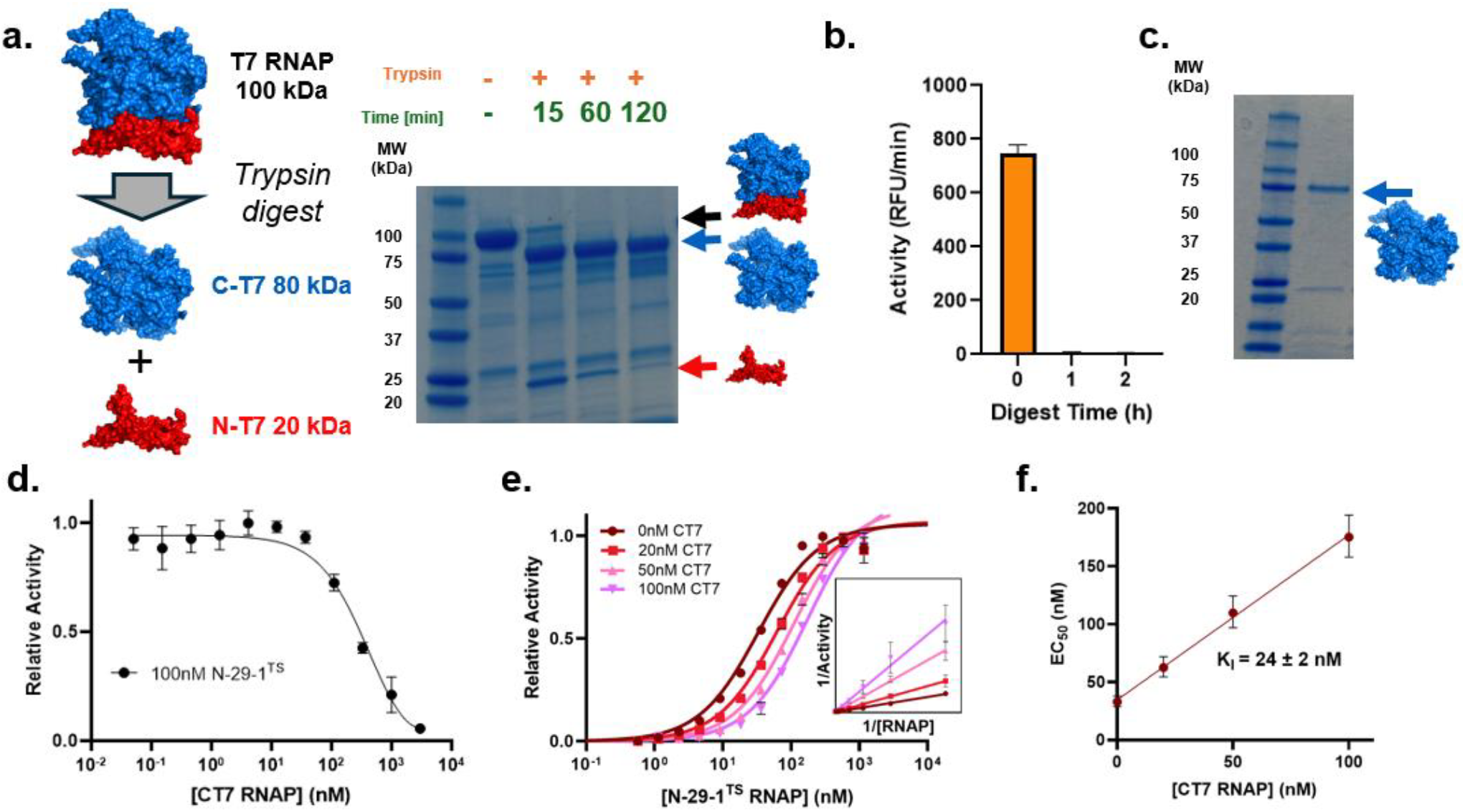
CT7 acts as a competitive inhibitor in in vitro T7 RNAP transcription. a. Proteolytic profile of 3 mg/mL N-29-1TS by trypsin digestion at a 750x mass excess of N-29-1TS at room temperature. b. Impact of protease digestion time on transcriptional activity for the N-29-1TS construct. c. CT7 purity after cation exchange chromatography. d. Relative activity of 100 nM N–29-1TS vs [CT7] using 3 nM template DNA and 1 μM HBC530. e. Relative activity vs. N-29-1TS concentration using 3 nM template DNA, 1 μM HBC530, with 0, 20, 50, and 100 nM CT7. e inset. 1/activity vs 1/[RNAP] with 0, 20, 50, and 100 nM [CT7] shown as a Lineweaver-Burk plot. f. EC50 of N-29-1TS vs CT7 concentration. K_I_ was determined from fit linear regression using eqn. 6 (R2 = 0.99). For panels b., d., e., and f., the error bars represent 1 s.d. of at least three technical replicates. In some cases, the error bars are smaller than the symbol.

To test whether CT7 inhibits T7 RNAP in the aptamer assay, CT7 was titrated with constant 100 nM N-29-1^TS^. CT7 was found to decrease the activity by half at equimolar ratios to the full-length N-29-1^TS^, and almost completely inhibit activity in gross excess (**Fig 2e**). To test our proposed competitive inhibition model, CT7 was added at various concentrations when titrating N-29-1^TS^. We found the EC_50_ of transcriptional activity increased with the addition of increasing CT7 (**Fig 2f**). Plotting the same data on a Lineweaver-Burk plot showed the same Y-intercept and different slopes with different CT7 concentrations, both hallmarks of a competitive inhibition (**Fig 2e**, inset). Indeed, the inferred EC_50_ shift was linearly related to the concentration of CT7 (**Fig 2g**). Fitting our proposed inhibition model results in an inhibition constant K_I_= 24 nM (22-26 95% c.i.). Thus, CT7 competitively inhibits T7 RNAP transcriptional activity with an inhibition constant approximately equivalent to the EC_50_ of the full-length N-29-1^TS^ construct.

## Discussion

In this work, we found the EC_50_ transcriptional activity of WT and TS T7 RNAP were in the range of 100-600 nM, while N-29-1^TS^ showed an EC_50_ up to an order of magnitude lower. We theorize the mutational profile of N-29-1^TS^ alters the ability of the polymerase to proceed from promoter binding to forming a stable elongation complex. These effects both modify the EC_50_ and lower the overall transcriptional activity. Mutational perturbation of the N-terminus of T7 RNAP has been shown to alter the abortive transcript cycle and RNA product purity profile^1^. Whatever the underlying mechanism, full-length RNA polymerases can be engineered for lower EC_50_ values. Choosing a more responsive polymerase, like N-29-1^TS^, may be important in the development of more sensitive biosensors. Whether a reconstituted split N-29-1 also has a similar EC_50_ value remains to be determined.

Akama^26^ previously used a similar transcriptional assay to measure T7 RNAP performance under different conditions like template DNA concentration. While they did not directly measure EC_50_ values, retrospective analysis of their WT activity profiles as a function of DNA concentration yields a similar EC_50_ to ours. However, our EC_50_ measurements are over an order of magnitude higher than a previous study testing T7 RNAP activity^27^. In that study, T7 RNAP is held constant, which results in a smaller EC_50_ than equation 6 because the T7 RNAP mass balance needs to account for polymerase actively transcribing the DNA. Although we found that our EC_50_ values were largely insensitive to free magnesium or other salt concentrations, other subtle differences in the assay conditions may also be reflected on the EC_50_ differences between studies. For now, which values to use for modeling *in vivo* T7-based genetic circuits ^28–30^ may depend on whether one is in a DNA-limiting vs. polymerase-limiting regime.

High EC_50_ values observed for split T7 RNAP biosensors may relate to protein translation, folding, and turnover processes *in vivo*. Alternatively, genetic circuits *in vivo* may not operate at the steady state assumed by the model. Nevertheless, our data supports CT7 as a competitive inhibitor for transcription by T7 RNAP. Since a single molecule is approximately 1 nM intracellular concentration in bacteria, a relatively small surplus of CT7 may lead to large changes in the responsiveness of biosensors using split T7 transduction. Thus, NT7 is strictly required in excess^31^. Our findings can explain some of the reasons why the biosensor EC_50_s are much higher than intrinsically expected by the chemically induced proximity partners. Even with no excess CT7, our findings support a theoretically minimal EC_50_ for N-29-1 -based biosensors.

## Conclusion

In conclusion, we adapted a biochemical network model and determined likely biochemical parameters for T7 RNAP and the inhibitory constant for CT7. Our results should be useful in predictive modeling of T7 RNAP genetic circuits.

## Acknowledgements

We thank B. Garcea, B. Dickinson, S. Cutler, and I. Wheeldon for helpful discussions related to production of CT7 and split biosensors. HBC530 was a kind gift from J. Yesselman.

## Author Contributions

Conceptualization: R.D, Z.T.B., T.A.W.; Designed bench research: R.D., K.A.L., Z.T.B., T.A.W.; Performed bench research: R.D., K.A.L., Z.T.B.; Developed model: O.I.I., T.A.W., Data analysis: R.D., K.A.L., Z.T.B., T.A.W.; Writing: R.D., T.A.W. with input from all co-authors; Supervision: T.A.W.; Funding Acquisition: Z.T.B, T.A.W.

## Funding Sources

This work was supported by the National Science Foundation (Award #: 2218330), the National Science Foundation Graduate Research Fellowship Program (Z.T.B. DGE Award Number 2040434, fellow ID: 2021324468). K.L was supported by the University of Colorado Boulder through participation in its Young Scholars Summer Research Program, with support from the National Science Foundation (Award #EEC-2348856) and the CU Department of Chemical and Biological Engineering.

## Notes

https://doi.org/10.5281/zenodo.13830256

### Competing Interest Statement

Timothy A. Whitehead reports a relationship with Metaphore Biotechnologies, Inc. that includes: board membership. Timothy A. Whitehead reports a relationship with Alta Resource Technologies, Inc. that includes: board membership. If there are other authors, they declare that they have no known competing financial interests or personal relationships that could have appeared to influence the work reported in this paper.

## References

1. Dousis, A., Ravichandran, K., Hobert, E. M., Moore, M. J. & Rabideau, A. E. An engineered T7 RNA polymerase that produces mRNA free of immunostimulatory byproducts. Nat. Biotechnol. 41, 560–568 (2023).

2. Kar, S. & Ellington, A. D. Construction of synthetic T7 RNA polymerase expression systems. Methods 143, 110–120 (2018).

3. Shis, D. L. & Bennett, M. R. Library of synthetic transcriptional AND gates built with split T7 RNA polymerase mutants. Proc. Natl. Acad. Sci. U. S. A. 110, 5028–5033 (2013).

4. Schaerli, Y., Gili, M. & Isalan, M. A split intein T7 RNA polymerase for transcriptional AND-logic. Nucleic Acids Res. 42, 12322–12328 (2014).

5. Chee, W. K. D., Yeoh, J. W., Dao, V. L. & Poh, C. L. Highly Reversible Tunable Thermal-Repressible Split-T7 RNA Polymerases (Thermal-T7RNAPs) for Dynamic Gene Regulation. ACS Synth. Biol. 11, 921–937 (2022).

6. Segall-Shapiro, T. H., Meyer, A. J., Ellington, A. D., Sontag, E. D. & Voigt, C. A. A ‘resource allocator’ for transcription based on a highly fragmented T7 RNA polymerase. Mol. Syst. Biol. 10, 742 (2014).

7. Baumer, Z. T. et al. Dynamic regulation of engineered T7 RNA polymerases by endogenous metabolites. 2024.08.07.607023 Preprint at 10.1101/2024.08.07.607023 (2024).

8. Schaffter, S. W. & Schulman, R. Building in vitro transcriptional regulatory networks by successively integrating multiple functional circuit modules. Nat. Chem. 11, 829–838 (2019).

9. Pu, J., Zinkus-Boltz, J. & Dickinson, B. C. Evolution of a split RNA polymerase as a versatile biosensor platform. Nat. Chem. Biol. 13, 432–438 (2017).

10. Pu, J., Disare, M. & Dickinson, B. C. Evolution of C-terminal modification tolerance in full-length and split T7 RNA Polymerase biosensors. Chembiochem Eur. J. Chem. Biol. 20, 1547–1553 (2019).

11. Park, S.-Y. et al. An orthogonalized PYR1-based CID module with reprogrammable ligand-binding specificity. Nat. Chem. Biol. 20, 103–110 (2024).

12. Komatsu, S., Ohno, H. & Saito, H. Target-dependent RNA polymerase as universal platform for gene expression control in response to intracellular molecules. Nat. Commun. 14, 7256 (2023).

13. Martin, N. T. et al. Engineering Rapalog-Inducible Genetic Switches Based on Split-T7 Polymerase to Regulate Oncolytic Virus-Driven Production of Tumour-Localized IL-12 for Anti-Cancer Immunotherapy. Pharm. Basel Switz. 16, 709 (2023).

14. McSweeney, M. A. et al. A modular cell-free protein biosensor platform using split T7 RNA polymerase. BioRxiv Prepr. Serv. Biol. 2024.07.19.604303 (2024) doi:10.1101/2024.07.19.604303.

15. Yuan, Y. & Miao, J. Agrochemical control of gene expression using evolved split RNA polymerase. PeerJ 10, e13619 (2022).

16. Jones, K. A., Snodgrass, H. M., Belsare, K., Dickinson, B. C. & Lewis, J. C. Phage-Assisted Continuous Evolution and Selection of Enzymes for Chemical Synthesis. ACS Cent. Sci. 7, 1581–1590 (2021).

17. Stanton, B. Z., Chory, E. J. & Crabtree, G. R. Chemically induced proximity in biology and medicine. Science 359, eaao5902 (2018).

18. Park, S.-Y. et al. Agrochemical control of plant water use using engineered abscisic acid receptors. Nature 520, 545–548 (2015).

19. Beltrán, J. et al. Rapid biosensor development using plant hormone receptors as reprogrammable scaffolds. Nat. Biotechnol. 40, 1855–1861 (2022).

20. Bayle, J. H. et al. Rapamycin Analogs with Differential Binding Specificity Permit Orthogonal Control of Protein Activity. Chem. Biol. 13, 99–107 (2006).

21. Ikeda, R. A. & Richardson, C. C. Enzymatic properties of a proteolytically nicked RNA polymerase of bacteriophage T7. J. Biol. Chem. 262, 3790–3799 (1987).

22. Muller, D. K., Martin, C. T. & Coleman, J. E. Processivity of proteolytically modified forms of T7 RNA polymerase. Biochemistry 27, 5763–5771 (1988).

23. Sugiyama, A., Nishiya, Y. & Kawakami, B. RNA polymerase mutants with increased thermostability. (2009).

24. Chen, X. et al. Visualizing RNA dynamics in live cells with bright and stable fluorescent RNAs. Nat. Biotechnol. 37, 1287–1293 (2019).

25. Chamberlin, M. & Ring, J. Characterization of T7-specific ribonucleic acid polymerase. General properties of the enzymatic reaction and the template specificity of the enzyme. J. Biol. Chem. 248, 2235–2244 (1973).

26. Akama, S., Yamamura, M. & Kigawa, T. A Multiphysics Model of In Vitro Transcription Coupling Enzymatic Reaction and Precipitation Formation. Biophys. J. 102, 221–230 (2012).

27. Arnold, S. et al. Kinetic modeling and simulation of in vitro transcription by phage T7 RNA polymerase. Biotechnol. Bioeng. 72, 548–561 (2001).

28. Fabrini, G. et al. Co-transcriptional production of programmable RNA condensates and synthetic organelles. Nat. Nanotechnol. 1–9 (2024) doi:10.1038/s41565-024-01726-x.

29. Tian, Z. et al. Circular single-stranded DNA as a programmable vector for gene regulation in cell-free protein expression systems. Nat. Commun. 15, 4635 (2024).

30. Singhal, V., Tuza, Z. A., Sun, Z. Z. & Murray, R. M. A MATLAB toolbox for modeling genetic circuits in cell-free systems. Synth. Biol. Oxf. Engl. 6, ysab007 (2021).

31. Steiner, P. J. et al. A Closed Form Model for Molecular Ratchet-Type Chemically Induced Dimerization Modules. Biochemistry 62, 281–291 (2023).

